# Methanolic extracts of the edible halophyte *Crithmum maritimum* enhance oxidative stress resistance in *Caenorhabditis elegans* through hormetic mechanisms

**DOI:** 10.1101/2023.07.19.549636

**Authors:** Raquel Martins-Noguerol, Alejandro Mata-Cabana, María Olmedo, Cristina DeAndrés-Gil, Xoaquín Moreira, Marta Francisco, Antonio J. Moreno-Pérez, Jesús Cambrollé

## Abstract

Halophytes are promising sources of bioactive phenolic compounds for the food and pharmaceutical industries. However, their phenolic composition is influenced by environmental conditions, and the *in vivo* antioxidant activity of their phytochemicals is largely unknown. We evaluated the antioxidant capacity of phenolic-rich methanolic extracts from the edible halophyte *Crithmum maritimum*, grown in wild and greenhouse conditions. Additionally, their *in vivo* antioxidant capacity was analyzed for the first time using the model *Caenorhabditis elegans*. Wild plant extracts showed higher phenolic content and diversity, and *in vitro* antioxidant activity. Both extracts enhanced oxidative stress resistance and increased nematode survival rates, albeit to varying extents, and increased reactive oxygen species production in nematodes, without affecting their lifespan, suggesting a hormetic mechanism. Although no neuroprotective effects were observed in models of neurodegenerative diseases, these findings highlight the potential of *C. maritimum* as a valuable source of phenolics with antioxidant properties for the food industry.

## 1. Introduction

Halophytes are salt-tolerant plants which have been traditionally consumed for their organoleptic and medicinal properties (Barreira et al., 2017). Throughout its evolution, halophytes acquired powerful antioxidant systems, such as the biosynthesis of phenolic compounds (e.g., flavonoids, tannins), to cope with the high environmental pressures where they naturally grow, particularly high levels of salinity (Ksouri et al., 2012). This makes halophytes an interesting source of dietary phytochemicals with bioactive properties (Oueslati et al., 2012; Lopes et al., 2021). Dietary polyphenols can minimize the deleterious effects of oxidative stress and exhibit anti-inflammatory activity, thereby promoting an optimal aging (Balasundram et al., 2006). Furthermore, current evidence supports an important role of these dietary antioxidants in the development/prevention of neurodegenerative disorders, such as Alzheimer’s and Parkinson’s disease (Costa et al., 2017).

Nevertheless, the total phenolic content and composition of halophytes are highly influenced by factors such as abiotic conditions and phenology (Ksouri et al., 2012; Mekinić et al., 2018). Additionally, antioxidant compounds are often selected based on their *in vitro* antioxidant activity, with the assumption that they will also exhibit antioxidant effects in living organisms. However, studies have shown that there is not always a correlation between both *in vitro* and *in vivo* effects (Pun et al., 2010; Mota et al., 2023). Additionally, some halophytes exhibit significant antioxidant capacity despite their comparatively lower total phenolic content, which can be attributed to the distinct activity of polyphenols inherent in their chemical composition (Lopes et al., 2021; Chen et al., 2020). Otherwise, some phenolic compounds have shown beneficial effects, while others have not or even showed potential harmful effects at high doses (Ristow, 2014; Alì et al., 2021). Therefore, it remains unclear to what extent these phytochemicals of halophytes contribute to the antioxidant *in vivo* effects, which is essential information to fully understand their functional value for commercial uses (Mota et al., 2023).

*Crithmum maritimum* L. (Apiaceae), known as sea fennel, is a wild edible and medicinal halophyte that grows in coastal areas throughout Western Europe and is highly valued for its nutritional and antioxidant properties (Castillo et al., 2022). However, its potential as a source of antioxidant compounds is based on studies about its phenolic content and *in vitro* assays, which may be greatly affected by abiotic factors (e.g., Meot-Duros & Magné, 2009; Mekinić et al., 2018). Furthermore, the *in vivo* antioxidant activity of its phytochemicals is currently unknown. In the present study, we evaluated the antioxidant capacity of phenolic-rich methanolic extracts from *C. maritimum* grown under different conditions (i.e., wild and optimal greenhouse conditions). We analyzed (i) the phenolic content and composition of methanolic extracts of *C. maritimum* leaves and their *in vitro* antioxidant activity by biochemical techniques; (ii) their *in vivo* antioxidant activity in the model organism *Caenorhabditis elegans*, and (iii) their potential neuroprotective effects using mutant lines of *C. elegans*. This is the first comprehensive study that evaluates the *in vivo* antioxidant properties of phenolics in an edible halophyte and how growing conditions can influence them.

## 2. Materials and Methods

### 2.1. Plant material and extract preparation

In mid-September 2020, we collected seeds from *C. maritimum* plants at Roche Beach (Spain, 36.314138/-6.153952) and sow them in plastic pots (n=6) filled with perlite in a greenhouse with natural daylight (200-1000 μmol m^-2^ s^-1^), 22-25 °C and 40-60% of relative humidity. Plants were watered with 20% Hoagland’s solution for optimal nutrient supply (Castillo et al., 2022). In spring 2022, we collected leaves from wild *C. maritimum* plants at Roche Beach and from the greenhouse plants. The extracts were obtained by grinding 1 g of fresh leaves with liquid nitrogen and adding 5 mL of 70% methanol. The mixture was ground on ice and incubated for 15 min, with shaking every 2 min followed by centrifugation. The supernatant was transferred to clean tubes and the extracts were kept at -20 °C.

### 2.2. Determination of phenolic compounds

Phenolic compounds were extracted from 20 mg of ground material with 0.25 mL of 70% methanol in an ultrasonic bath for 15 min, followed by centrifugation. The supernatant was filtered (0.20-µm micropore PTFE membrane) and placed in vials for chromatographic analysis. For chemical identification of the polyphenols, we used ultra-performance liquid chromatography coupled with an electrospray ionization quadrupole (Thermo Dionex Ultimate 3000 LC) time-of-flight mass spectrometry (UPLC-Q-TOF-MS/MS; Bruker Daltonics GmbH) with a heated electrospray ionization (ESI) source, according to Castillo et al. (2022). Polyphenols were identified based on the data obtained from the standard substances or published literature including RT, λmax, ([M–H]^−^), and major fragment ions. For quantification of phenolic compounds, 10 µL of each sample was analyzed using the same column and conditions used for identification, in an UHPLC (Nexera LC-30AD; Shimadzu Corporation) equipped with a Nexera SIL-30AC injector and one SPD-M20A UV/VIS photodiode array detector (Moreira et al., 2021). The flavonoids were quantified as rutin equivalents and phenolic acids as chlorogenic acid equivalents by external calibration using calibration curves.

### 2.3. DPPH radical scavenging assay

The *in vitro* antioxidant activity of the extracts was evaluated according to Coppari et al. (2021). 50 µL of plant extract dilutions (from 1/5 to 1/100) were mixed with 150 µL of a methanolic solution of DPPH (2,2-diphenyl-1-picryl-hydrazyl-hydrate; Sigma-Aldrich) in a 96-well plate. The absorbance was measured at 517 nm. DPPH inhibition (I) was calculated as follows:

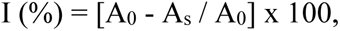

where A_0_ represents the absorbance of the negative control and A_s_ is the absorbance of the sample. We used 70% methanol as the blank and butylated hydroxytoluene (Sigma-Aldrich) as a standard control. The EC_50_ value was determined using the equation derived from the linear regression analysis.

### 2.4. *Caenorhabditis elegans* strains and maintenance

All strains used in this study were obtained from the Caenorhabditis Genetics Center. N2 Bristol strain was used as wild type. The strain GMC101 (*dvIs100* [*unc-54p::A-beta-1-42::unc-54* 3′-UTR + *mtl-2p:*:GFP]) was used as model for Alzheimer’s disease (McColl et al., 2012). The strain AM141 (*rmIs133* [*unc-54p*::Q40::YFP]) was used as a model for polyglutamine diseases (Huntington’s disease; Morley et al., 2002). Nematodes were cultured and maintained following standard methods (Brenner, 1974), typically at 20 °C on Nematode Growth Medium (NGM) plates supplemented with *Escherichia coli OP50-1* as a food source (Mata-Cabana et al., 2022). *In vivo* assays were conducted using a non-lethal submaximal concentration of the extracts (1/10 dilution in 70% methanol) and 70% methanol was used as control treatment.

### 2.5. Caenorhabditis elegans lifespan assay

Lifespan was assessed following standard methods (Lionaki & Tavernarakis, 2013). Gravid adults were treated with an alkaline hypochlorite solution to obtain embryos, which were then placed in NGM plates with *E. coli* OP50-1, and 500 µL of either *C. maritimum* extracts or 70% methanol spread across the entire surface. Then, 50 synchronized L4 stage animals were transferred to new NGM plates with the treatments. Nematodes were transferred to new plates every 2 days until reproduction ceased. Live and dead (unresponsive to prodding with a platinum wire) animals were daily counted. The entire assay was conducted at 20 °C. Survival curves were generated and analyzed using the GraphPad Prism software.

### 2.6. Oxidative stress resistance assay in *Caenorhabditis elegans*

Hydrogen peroxide (Sigma-Aldrich) was used to induce the oxidative stress. Embryos were obtained from gravid adults and placed in NGM plates with *E. coli OP50-1* and the treatments. L4 animals were transferred to new NGM plates with 1 mM hydrogen peroxide. The assay was conducted at 20 °C for 3 hours, and the number of living animals was scored. Animals unresponsive to prodding with a platinum wire were considered dead. The assay was repeated three times (50 animals/trial). The final percentage of living animals was calculated.

### 2.7. Quantification of intracellular ROS in *Caenorhabditis elegans*

Reactive oxygen species (ROS) quantification was performed in 10-days adult nematodes. Nematodes were collected in M9 buffer supplemented with 0.01% polyethylene glycol, washed twice, and transferred to tubes with M9 buffer. Nematodes were incubated for 2 h at 20 °C with 10 μM of dihydroethidium (DHE, Sigma). Then, animals were anesthetized with 20 mM Levamisole and red fluorescent signal of DHE was imaged and recorded in a Leica M205 FCA stereo microscope. The intensity of the signal was later quantified using the Image J software. DHE signal was measured in at least 15 animals per condition for 3 biological replicates.

### 2.8. Antioxidant effects in neurodegenerative disorders

We tested the effects of the extracts on nematode models for Alzheimer’s (GMC101) and Huntington’s (AM141) disease. Shifting GMC101 animals from 20 °C to 25 °C causes paralysis due to the Aβ_42_ aggregation (McColl et al., 2012). L4 animals were tested for paralysis after 24 h hours at 25 °C by tapping their noses with a platinum wire. AM141 animals express a stretch of 40 glutamines (Q40) fused to the yellow fluorescent protein (YFP). When they reach adulthood, they show an entirely Q40::YFP aggregated phenotype (Morley et al., 2002). The aggregate formation was visualized using a Leica M205 FCA stereo microscope and the number of foci per animal was counted.

### 2.9. Statistical analyses

Statistical analyses were performed using IBM SPSS v. 24.0 software (IBM Corp.). Data were tested for normality (Kolmogorov-Smirnov test) and for homogeneity of variance (Levene test) and then analysed by t-Student and ANOVA. Tukey test was used as post hoc. For non-parametric data, Kruskal-Wallis test followed by Mann-Whitney U test were employed.

## 3. Results and Discussion

### 3.1. Phenolic composition and *in vitro* antioxidant activity

Wild *C. maritimum* plants showed a 60% higher total phenolic content (TPC) compared to greenhouse plants (Table 1). Both extracts contained phenolic acids and flavonoids, although the wild plants had higher proportion of flavonoids, accounting for 86.4% compared to 61.6% in greenhouse plants. The extracts from wild plants showed stronger *in vitro* antioxidant activity, as they achieved 50% DPPH inhibition (EC_50_) at a lower concentration (Table 1). This suggests that the wild plants, which likely experienced environmental stresses such as salinity and nutrient deprivation, have higher *in vitro* antioxidant activity, which correlated with TPC. Polyphenols are known to be produced as a response to extreme environmental conditions, in addition to genetic factors (Ksouri et al., 2012). Wild *C. maritimum* exhibited comparable or even higher TPC than other wild edible halophytes that are well-known sources of antioxidants, such as *Salicornia ramosissima* (33.0 mg/g DW) and *Sarcocornia perennis* (20.5 mg/g DW) (Barreira et al., 2017). Otherwise, the TPC of greenhouse plants (Table 1) was higher than other wild edible halophytes like *Mesembryanthemum nodiflorum* or *Sarcocornia fruticosa* (5.12-6.81 mg/g DW; Castañeda-Loaiza et al., 2020).

**Table 1.** Total phenolic content (TPC; mg/g dry weight, DW), flavonoids and phenolic acids (%) and DPPH scavenging activity (EC_50_; μg/mL) of methanolic extracts of wild and greenhouse plants of *C. maritimum*. Data represent average and standard error of twelve (wild) and three (greenhouse) independent replicates. (***) Statistical differences between methanolic extracts (p<0.001).

|  | **TPC** | **Flavonoids** | **Phenolic acids** | **DPPH** |
| --- | --- | --- | --- | --- |
| Wild | 30.3 ± 1.0 | 86.4 ± 0.8 | 13.6 ± 0.8 | 1.3 ± 0.1 |
| Greenhouse | 12.7 ± 1.0 *** | 61.6 ± 4.3*** | 38.4 ± 4.3*** | 16.1 ± 0.0*** |

Additionally, significant differences were observed in the phenolic composition of the plant extracts (Figure 1). Wild plants showed a dominant flavonoid, rutin (56-75% of TPC), along with two quercetin-type compounds (9-31% of TPC), while greenhouse plants contained kaempferol 3-glucoside-7-rhamnoside as the sole flavonoid (57-70% of TPC; Figure 1A). Regarding phenolic acids, ferulic acid was exclusively detected in wild plants, while the proportion of different caffeoylquinic acid isomers (chlorogenic acid) was higher in greenhouse plants (Figure 1B). Therefore, the distinct chemical profiles of the extracts could also contribute to their antioxidant activity (Lopes et al., 2021; Chen et al., 2020). Previous studies have shown that phenolic parameters can vary within the same species due to multiple factors affecting accumulation and degradation processes (Bertin et al., 2014), which support our results. These findings indicate that *C. maritimum* has the potential to be a valuable source of phenolics with antioxidant activity, which could be influenced by growing conditions. However, the correlation between *in vitro* and *in vivo* effects may vary (Mota et al., 2023), highlighting the importance of *in vivo* assays to understand their antioxidant properties.

**Figure 1.**
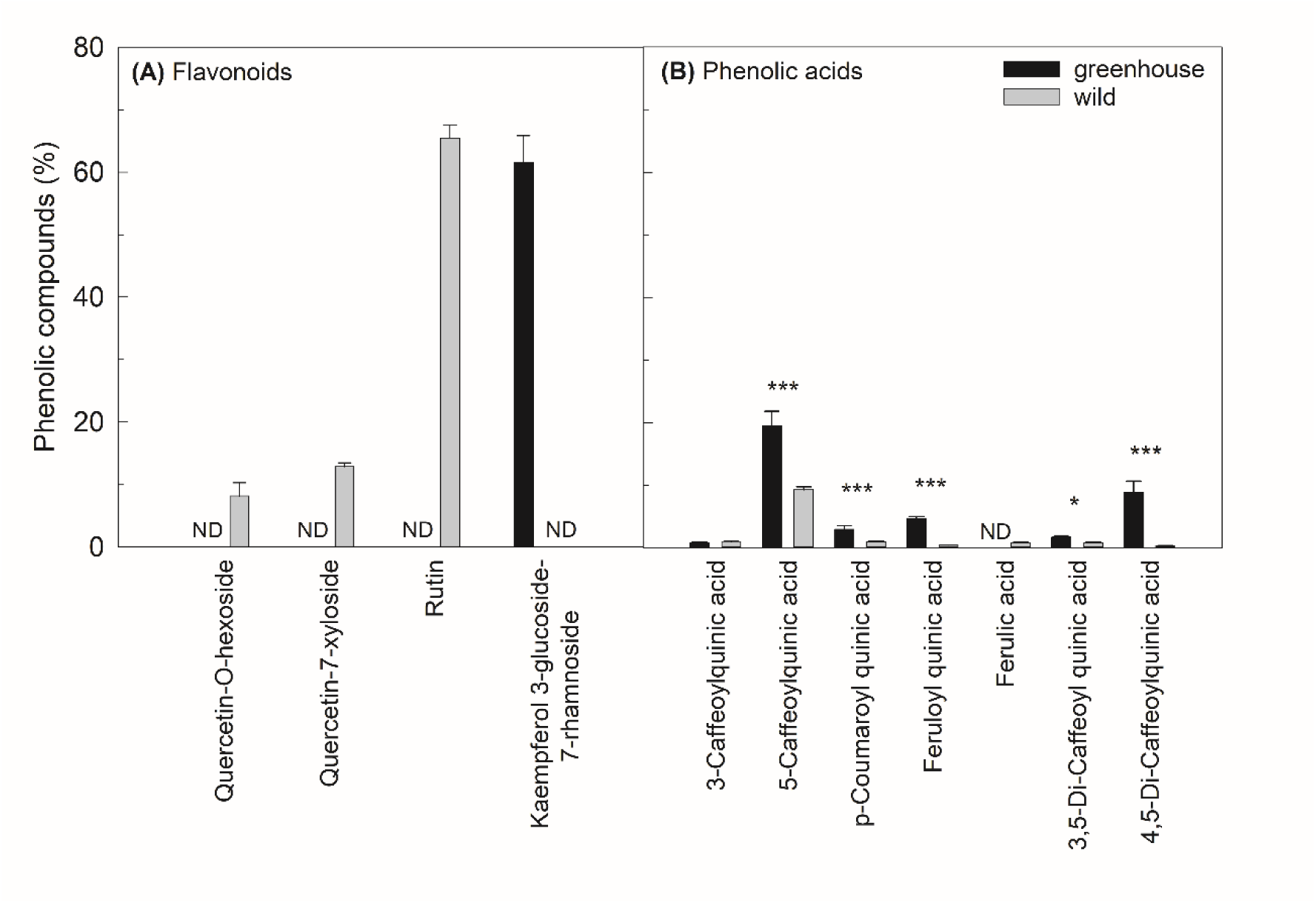
Phenolic compounds (%) in *C. maritimum* leaves from greenhouse and wild plants. **(A)** Flavonoids. **(B)** Phenolic acids. Data represent average and standard error of three (greenhouse) and twelve (wild) independent replicates. Asterisks represent statistical differences: (*) p<0.05, (**) p<0.01, (***) p<0.001. ND, non detected.

### 3.2. *In vivo* antioxidant activity

The *in vivo* antioxidant effects of *C. maritimum* extracts were evaluated in the nematode *C. elegans* as a model. When exposed to pro-oxidant compounds, such as hydrogen peroxide, *C. elegans* experiences oxidative stress, resulting in elevated levels of ROS and shortening the nematode survival (Gems & Doonan, 2009). After treatment with methanolic extracts, the nematodes significantly increased their survival to hydrogen peroxide-induced oxidative stress (Figure 2A). The extracts from wild plants showed a remarkable 40% increase in survival, while extracts from greenhouse plants showed an 18% increase, both compared to the control group. The extracts’ ability to neutralize free radicals may explain their response to oxidative stress. Rutin and quercetin, the main phenolics found in the extracts from wild *C. maritimum*, have been reported to increase stress resistance in nematodes (Kampkötter et al., 2007; Ayuda-Durán et al., 2019). Similarly, kaempferol, the dominant compound in the extracts from greenhouse plants, enhanced the survival of *C. elegans* under oxidative stress by modulating ROS accumulation (Kampkötter et al., 2007). These findings demonstrate the *in vivo* antioxidant activity of both extracts and their ability to enhance stress resistance in nematodes. Our results agree with previous studies in *C. elegans* with phenolic-rich plant extracts under oxidative stress conditions (González-Peña et al., 2021; Huang et al., 2022), and highlight the value of this model to screen halophyte phytochemicals with antioxidant properties.

**Figure 2.**
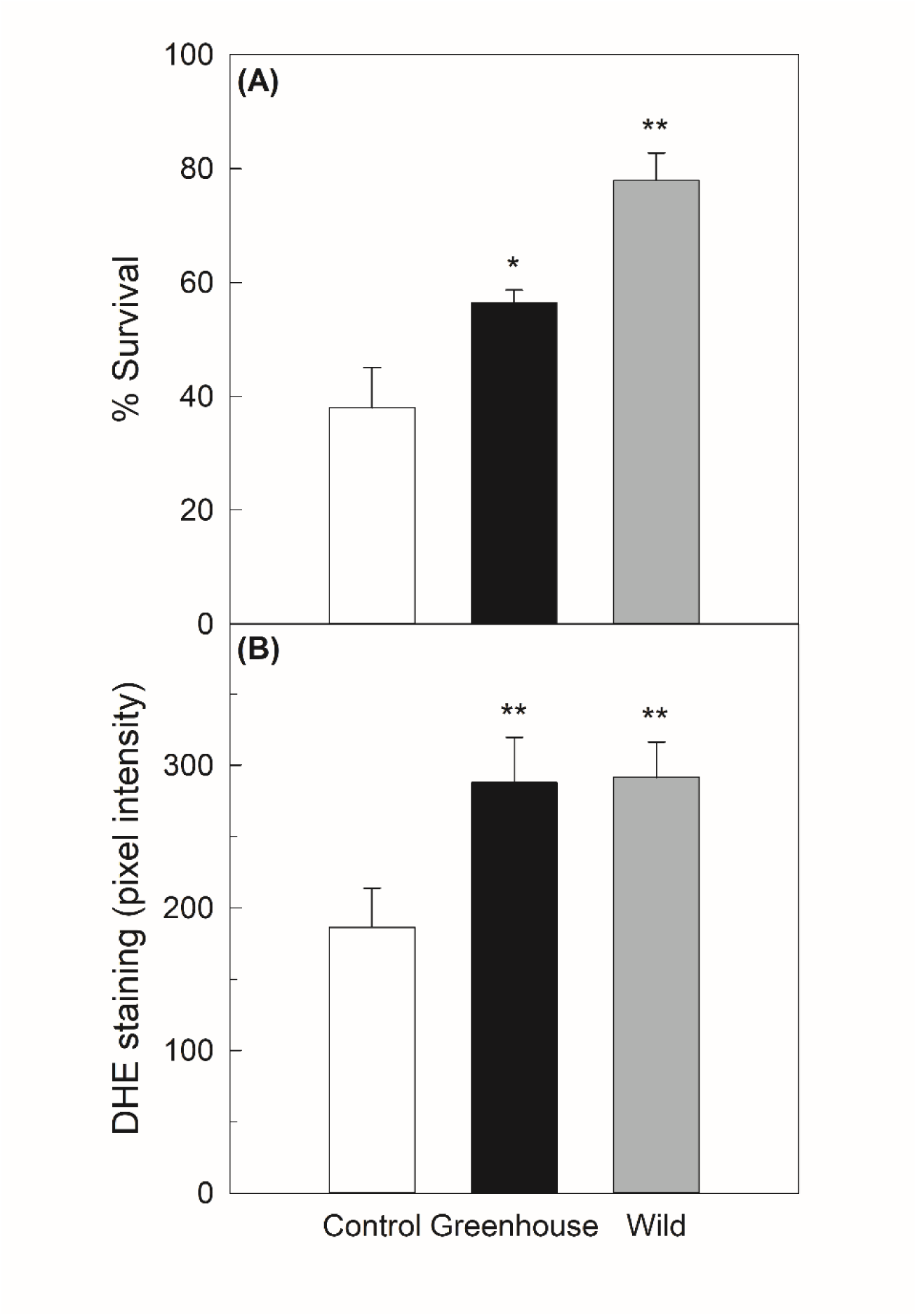
**(A)** Percent survival of *C. elegans* nematodes grown with *C. maritimum* extracts (from greenhouse or wild plants) after treatment with 1 mM hydrogen peroxide for 3 hours. **(B)** Quantification of ROS accumulation in the worm cultures after 10 days of treatment with *C. maritimum* extracts. Asterisks represent statistically significant differences: (*) p<0.05, (**) p<0.01.

On the other hand, animals treated with both extracts but not exposed to oxidative stress exhibited a higher intensity of DHE staining (Figure 2B), indicating an accumulation of ROS content compared to the control group under normal growth conditions. Previous studies have reported that plant extracts rich in antioxidants exert their protective effects by inducing a mild elevation of ROS that triggers cellular adaptive responses (Forman et al., 2014; Govindan et al., 2018; Li et al., 2019), which support our findings. Besides their radical scavenging ability, dietary antioxidants, such as plant polyphenols, are believed to act as pro-oxidants at low doses, activating mild oxidative stress and defense mechanisms to promote stress resistance (Ristow, 2014; Alì et al., 2021; González-Peña et al., 2021). This phenomenon, known as hormesis, suggests that high doses of exposure could be harmful, whereas low doses can activate protective pathways (Calabrese & Kozumbo, 2021). Compounds exhibiting hormetic effects typically enhance mitochondrial activity and ROS production, triggering adaptive stress response pathways (Li et al., 2019), enhancing an organism’s ability to cope with subsequent stress exposure (Ristow, 2014).

Otherwise, transient increases in intracellular ROS levels are known to play a crucial role in the aging and longevity of *C. elegans* (Miranda-Vizuete & Veal, 2017), with excessive ROS leading to a shortened lifespan (Zhu et al., 2022). Interestingly, none of the *C. maritimum* extracts influenced the lifespan of the animals (Figure 3). This is consistent with previous studies involving dietary polyphenols, such as the well-known resveratrol, which demonstrated protective effects against oxidative stress but did not extend the lifespan of *C. elegans* (Chen et al., 2013). Similar results were observed in nematodes treated with extracts from different plant species rich in antioxidant compounds (Pun et al., 2010). However, it should be noted that other plant extracts containing phenolics and isolated antioxidants have been shown to increase nematode lifespan (Govindan et al., 2018; Dilberger et al., 2021). Conflicting results in the literature could be due to the complexity of active compounds present in plant extracts, their chemical structure, concentrations, and interactions (Gallardo et al., 2006; Grünz et al., 2012).

**Figure 3.**
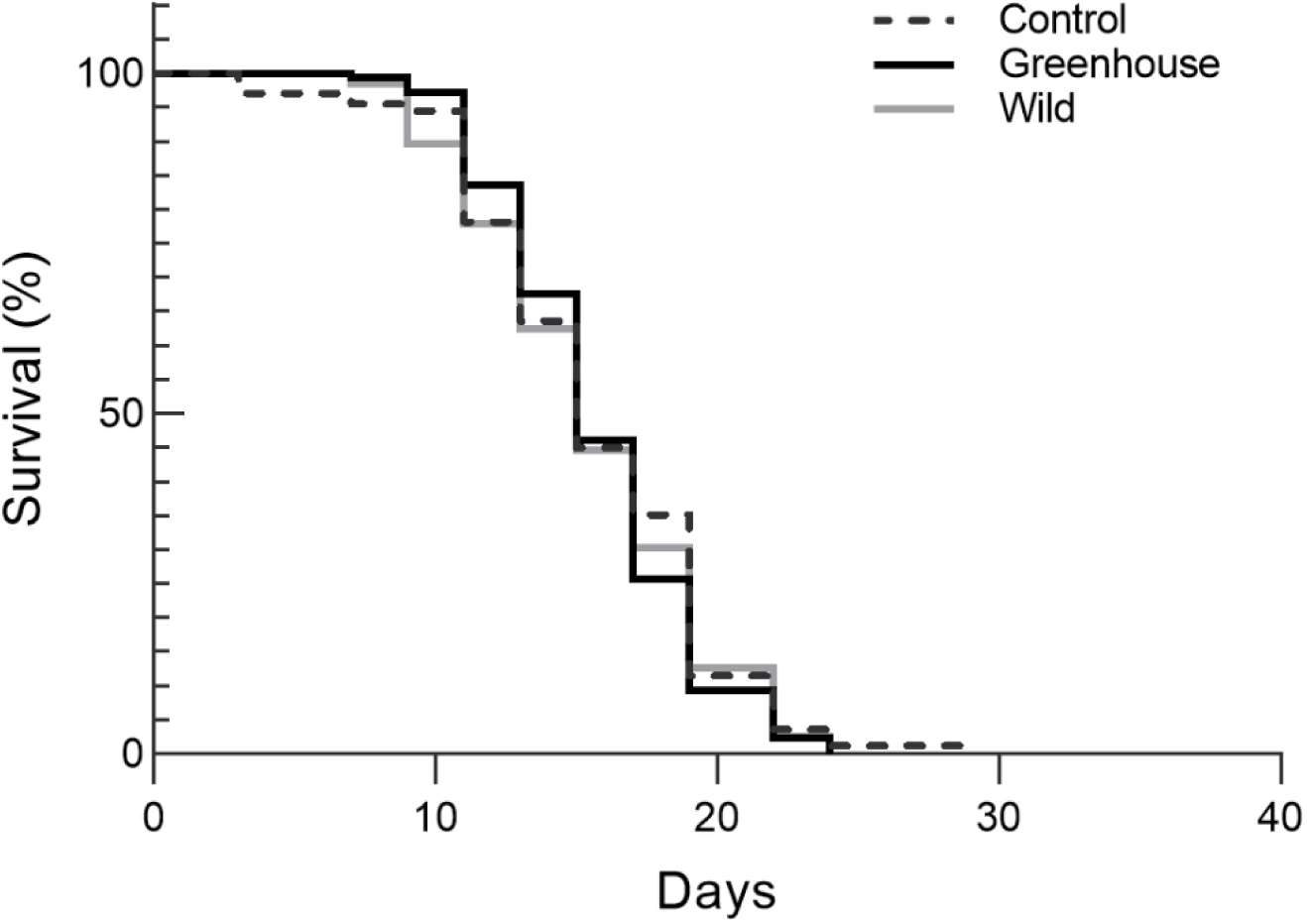
Survival curves of *C. elegans* nematodes grown with *C. maritimum* extracts from greenhouse or wild plants. Results are displayed as the percentage of nematodes that are alive at each time point. Each line represents the average of three independent replicates.

Our findings suggest that *C. maritimum* extracts have limited protective capacity in the absence of exogenous stress, but they exhibit a hormetic function by effectively safeguarding against oxidative stress, particularly the extract from wild plants. The increase in ROS levels may play a beneficial role by activating stress resistance, as previously reported (Miranda-Vizuete & Veal, 2017; Li et al., 2019), without affecting the lifespan of nematodes raised in optimal laboratory conditions. Further studies are required to comprehensively understand the cellular mechanisms underlying the hormetic response elicited by the phytochemicals present in *C. maritimum* extracts.

### 3.3. Neuroprotective effects

The intake of natural polyphenols has been associated with neuroprotective effects and their potential in preventing/treating neurodegenerative diseases (Costa et al., 2017). We investigated the neuroprotective effects of *C. maritimum* extracts using *C. elegans* models for Alzheimer’s and Huntington’s diseases, which induce paralysis and polyglutamine aggregates, respectively. However, the methanolic extracts did not reduce the percentage of paralyzed nematodes in the Alzheimer’s model or decrease polyQ aggregates in the Huntington’s model compared to the control groups (Table 2). These results indicate a lack of neuroprotective effects associated with the extracts. Previous studies suggest that chlorogenic acid in ethanolic *C. maritimum* extracts could be relevant in the treatment of neurodegenerative disorders (Mekinić et al., 2018). In our study, chlorogenic acid (caffeoyl quinic acid isomers) accounted for 11.4% and 30.8% of TPC in wild and greenhouse plants, respectively (equivalent to 3.4 and 3.9 mg/g DW), which is lower than the 16.3 mg/g DW content reported by Mekinić et al. (2018). The lower content of chlorogenic acid in the methanolic extracts, along with subsequent dilutions to avoid nematode toxicity, may explain these conflicting results. Nonetheless, it is worth noting that rutin and kaempferol, the primary phenolics in the methanolic extracts of wild and greenhouse plants have previously demonstrated protective effects against neurodegenerative pathologies in *C. elegans* (Sharoar et al., 2012; Cordeiro et al., 2020). Assessing the efficacy of plant extracts with different phenolic compositions can be challenging, as combinations of phytochemicals can yield contrasting effects (Costa et al., 2017). Further research is needed to unlock the unexplored potential of *C. maritimum* in addressing these diseases, utilizing diverse wild populations and greenhouse-grown plants under varying growing conditions.

**Table 2.** Percentage of paralyzed GMC101 nematodes (Alzheimer’s disease model) and number of protein aggregates in AM141 nematodes (Huntington’s disease) treated with methanolic extracts of *C. maritimum* and 70% methanol (control conditions). Data represent average and standard error of three independent replicates.

|  | **Percentage of paralyzed**<br>**GMC101 nematodes** | **Number of aggregates in**<br>**AM141nematodes** |
| --- | --- | --- |
| Control | 87.7 ± 5.8 | 68.0 ± 3.6 |
| Wild | 85.8 ± 6.1 | 70.7 ± 1.7 |
| Greenhouse | 86.2 ± 4.8 | 70.6 ± 0.0 |

## Conclusions

The methanolic extracts of wild *C. maritimum* plants exhibited higher phenolic content and antioxidant activity compared to greenhouse plants, with different phenolic composition. Both extracts demonstrated *in vivo* antioxidant effects and stress resistance in *C. elegans,* likely due to an increase in ROS accumulation underlying a hormetic response, but without affecting the nematode lifespan. Despite lacking neuroprotective effects in nematode models of neurodegenerative diseases, our findings highlight the significant potential of the edible halophyte *C. maritimum* as a valuable source of phenolic compounds with beneficial properties, making it a promising candidate for the food and nutraceutical industry.

## Acknowledgements

This work was financially supported by two grants from the Spanish Ministry of Science, Innovation and Universities (RTI2018-099260-A-I00 to J. Cambrollé and RTI2018-099322-B-100 to X. Moreira). R. Martins-Noguerol was financially supported by the Spanish Universities Ministry and European Union - Next Generation EU. We thank the Seville University Greenhouse General Service for their collaboration. A.J. Moreno-Pérez is the recipient of a research contract from the VII PPIT-US.

